# Splicing-aware resolution of scRNA-Seq data

**DOI:** 10.1101/2024.03.25.586675

**Authors:** D.K. Lukyanov, E.S. Egorov, V.V. Kriukova, K. Ladell, D. Price, A. Franke, D.M. Chudakov

## Abstract

Single-cell RNA sequencing (scRNA-Seq) provides invaluable insights in cell biology. Current scRNA-Seq analytic approaches do not distinguish between spliced and unspliced mRNA. RNA velocity paradigm suggests that the presence of unspliced mRNA reflects transitional cell states, informative for studies of dynamic processes such as embryogenesis or tissue regeneration. Alternatively, stable cell subsets may also maintain unspliced mRNA reservoirs for prompt initiation of transcription-independent expression. Based on the latter paradigm, we have developed a method called SANSARA (Splicing-Aware scrNa-Seq AppRoAch) for the splicing-aware analysis of scRNA-Seq data. We employed SANSARA to characterize peripheral blood regulatory T cell (T_reg_) subsets, revealing the complex interplay between FoxP3 and Helios master transcription factors and other unexpected splicing-informed features. For Th1 and cytotoxic CD4^+^ T cell subsets, SANSARA also revealed substantial splicing heterogeneity across crucial subset-specific genes. SANSARA is straightforward to implement in current data analysis pipelines and opens new dimensions in scRNA-Seq-based discoveries.

## Introduction

RNA processing is an integral part of the implementation of genetic information^1-2^. Correspondingly, rational utilization of splicing information in scRNA-Seq data analysis could reveal multiple functional aspects of cell biology. However, quantitative analysis of splicing is rarely included in scRNA-Seq studies due to the difficulties inherent to the short-read sequencing technologies^3-4^. Coverage bias across genes and sequencing technologies, inability to detect all splicing junctions, insufficient sequencing depth, and high dropout rate prevent direct estimation of splicing by distinguishing spliced and unspliced molecules^3^.

To date, splicing was studied in scRNA-Seq data in terms of transcriptional dynamics and cell-state transitions^5,6^, and only in a *post hoc* manner – after conventional clustering and dimensionality reduction. However, splicing information has not been used as an independent criterion to distinguish between stable functional cell subsets, implemented as an input at the level of cell clustering. At the same time, various cell subsets may preferentially accumulate unspliced primary transcripts in the nucleus, which can serve as transcription-independent reservoirs for rapid production of functional mature mRNA and proteins^7-8^. The same logic may be applicable to the non-coding RNA transcripts^9^. This means that one could consider the presence of certain unspliced RNA transcripts as a distinguishing feature for stable or relatively stable cell subsets, theoretically enabling the construction of splicing-aware scRNA-Seq data and the identification of corresponding functional cell clusters.

In this work, we describe SANSARA (Splicing-Aware scrNa-Seq AppRoAch), a method that produces splicing-adjusted gene expression matrix (saGEX) that accounts for the extent of splicing for each gene in each cell. The resulting saGEX is then subjected to a conventional clustering and dimensionality reduction pipeline to reconstruct a splicing-aware representation of the scRNA-Seq data.

We employ SANSARA to resolve the complexity of human peripheral blood helper T cells. This splicing-aware approach yields a deep structuring of the intrinsic heterogeneity of regulatory T cells (T_reg_s) and the Th1/cytotoxic axis of helper T cells. We anticipate that SANSARA should have broad applications in single-cell transcriptomics beyond T cell biology, revealing a universe of distinctive and informative splicing-related features of tissue cell subsets.

## Results

### Splitting gene expression into spliced and unspliced values

Direct estimation of the proportion of spliced versus unspliced mRNA for each gene in scRNA-Seq data is confounded by the oligo-dT primers used to enrich for polyadenylated mRNA molecules, and the limited coverage and biases of currently-available information obtained via either 5’-or 3’-high-throughput transcriptomics^3^. We settled on the veloVI framework^10^, which is based on the proportions of spliced and unspliced unique molecular identifiers (UMIs), where each UMI-labeled molecule containing a read mapping to an intronic region is counted as an unspliced molecule. These algorithms were initially developed for the determination of ‘RNA velocity’^6^, a parameter that reflects a transcriptomic snapshot of current mRNA turnover. Here we employed the veloVI-derived values to analyze cell heterogeneity using splicing-aware clustering and dimensionality reduction, in order to differentiate stable cell clusters characterized by distinct gene splicing features (**Fig. 1**).

**Figure 1.**
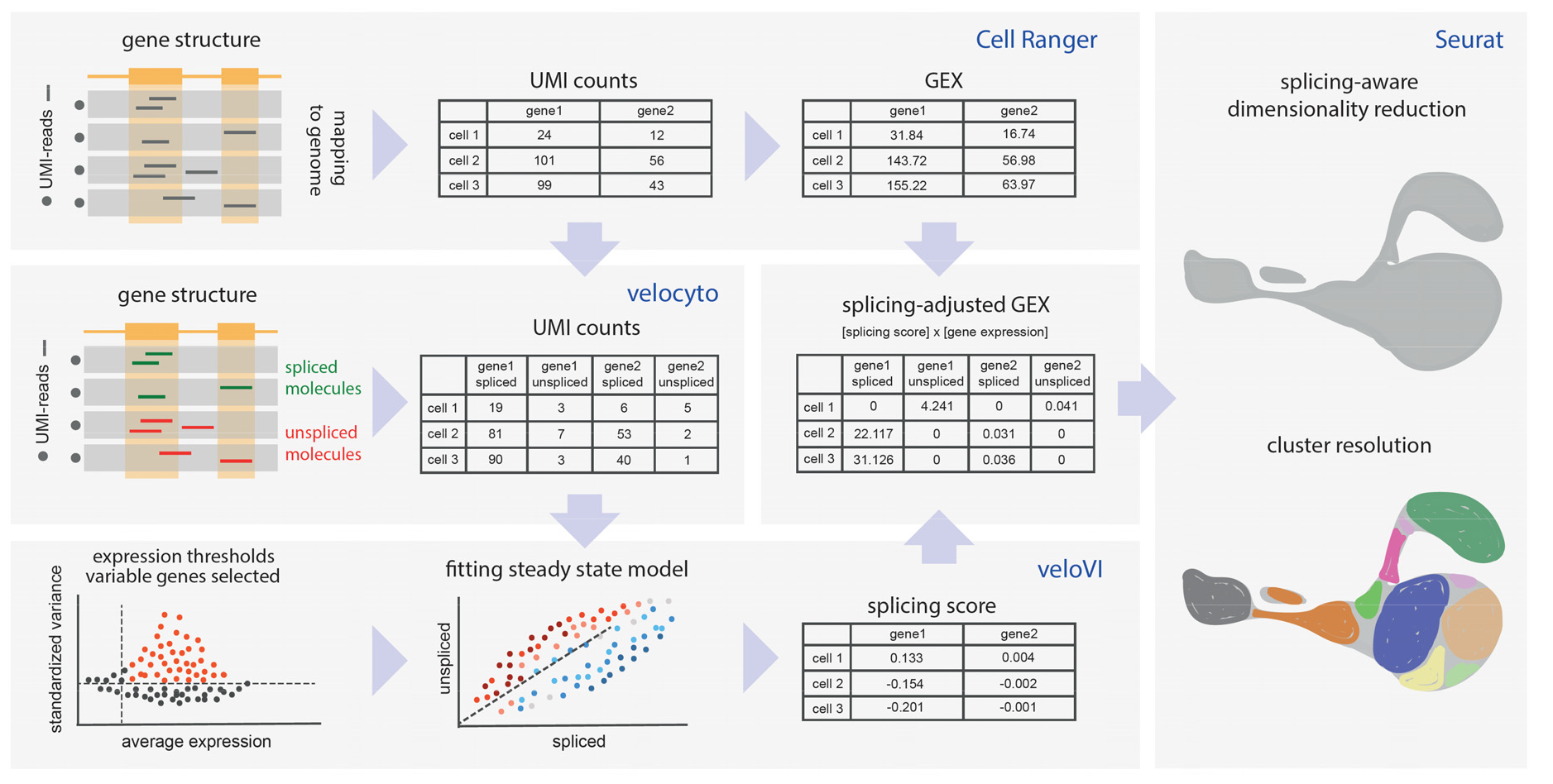
SANSARA workflow. After mapping scRNA-Seq data to the genome with Cell Ranger, spliced and unspliced UMI counts are differentiated using velocyto. Highly variable genes are selected based on log-normalized splicing-aware counts, and veloVI model is fitted to each gene. Genes for downstream analysis are chosen based on the quality of the fit. The product of original gene expression (GEX) and splicing score, termed splicing-adjusted GEX (saGEX) is then used for conventional dimensionality reduction and clustering analysis.

Initial velocyto-derived values^6^ depend on individual gene features and cannot be employed for informative analysis. The downstream veloVI-derived values represent a much more accurate individual estimation of the extent of gene splicing based on the inferred gene-specific rates of transcription, splicing, and degradation^10^. We used the veloVI filtering steps and confidence scores to choose the subset of genes most suitable for splicing estimation. Next, we used veloVI-derived values to calculate the splicing-adjusted gene expression (saGEX) for each gene. saGEX is determined by multiplying the veloVI value by the total normalized expression of that gene in each cell (GEX), and assigned to either the spliced (negative splicing score values) or the unspliced (positive splicing score values) form of the gene in each cell (**Fig. 1**).

The resulting saGEX cell-feature matrix simulates gene expression patterns conventionally used by dimensionality reduction methods, but split to discriminate spliced and unspliced gene forms. Finally, the saGEX data are analyzed using a standard Seurat pipeline—analogous to conventional GEX—with Seurat lognormalization. This approach, named SANSARA, proved to provide natural and informative downstream analyses, as demonstrated on the following examples.

### Resolving splicing differences in scRNA-Seq landscape

To test SANSARA, we used datasets of sorted, effector-enriched CD4^+^ T cells from peripheral blood mononuclear cells (PBMC) of three donors, which were extensively characterized previously^11^. Notably, these CD4^+^ 5’-RACE 10x Genomics scRNA-Seq datasets were of high quality, with more than 5,000 median UMIs per cell, sequenced with relatively long reads (100 + 100 nt) and high coverage of 90,000 reads per cell. This may be crucial for performance of the veloVI and SANSARA algorithms.

After obtaining saGEX values for ∼1,500 genes from individual donors, we integrated them using the Harmony pipeline^12^ to remove donor-specific batch effects (**Supplementary Fig. 1a,b**). The original splicing-unaware GEX datasets were analyzed separately using the same parameters for integration and dimensionality reduction. The general topology of the resulting splicing-aware UMAP data representation closely resembled that of the conventional splicing-unaware dataset, preserving the major subset composition (**Fig. 2a,b**). Splicing-aware UMAP plots were characterized by higher clustering stability at different resolutions (**Supplementary Fig. 1c–e**). Altogether, splicing-unaware and splicing-aware integration and clustering worked similarly, even though the splicing-aware dataset contains fundamentally different information.

**Figure 2.**
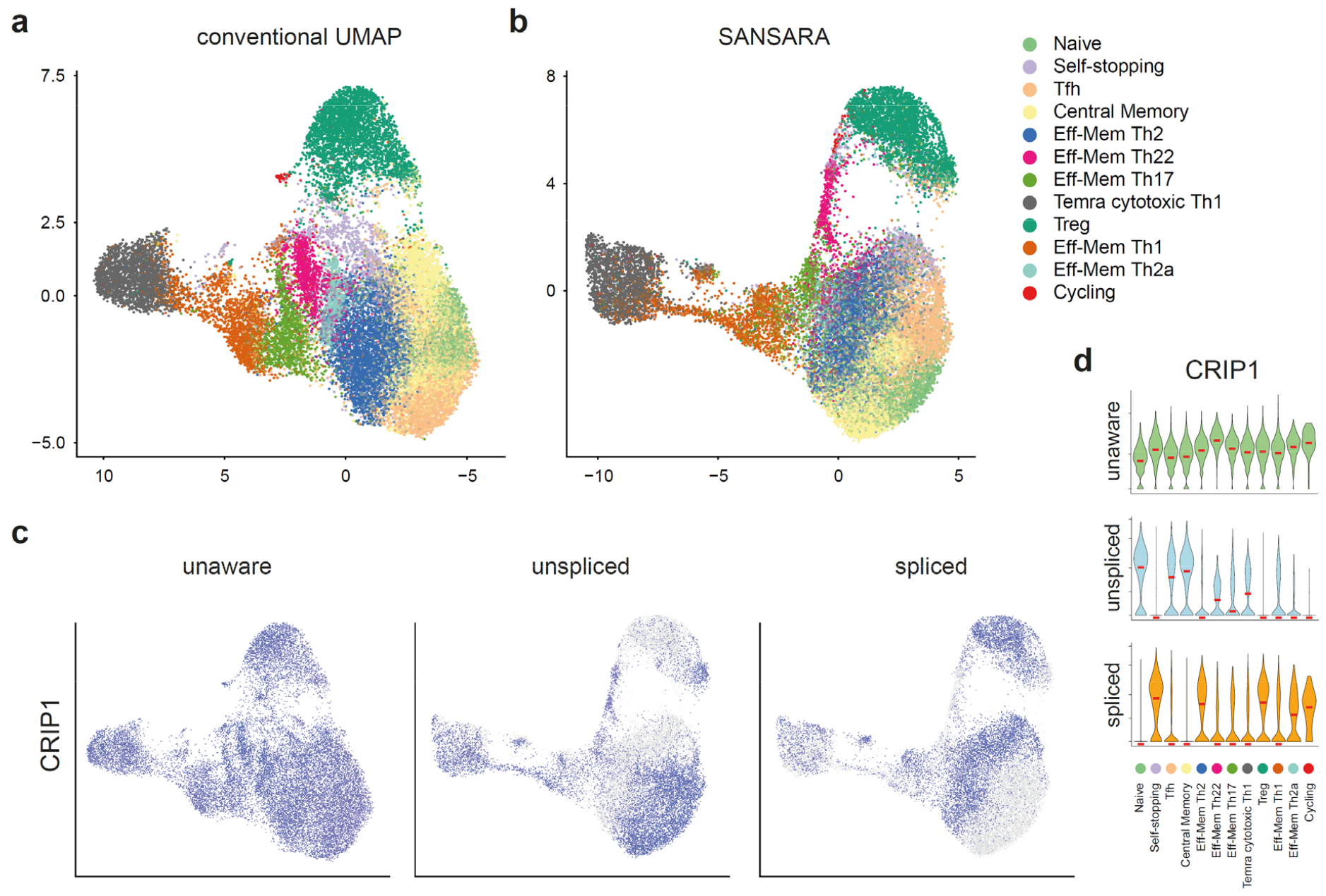
SANSARA reveals splicing heterogeneity of CD4^+^ T cells. **a,b**. Comparison of cluster annotation between the conventional splicing-unaware (a) and splicing-aware (b) UMAP plots. Annotated according to Ref. 11. **c**. UMAP plots of conventional GEX (left) versus saGEX (center, right) *CRIP1* expression. **d**. Violin plots of splicing-unaware (top) and -aware (middle, bottom) *CRIP1* expression across clusters.

Accounting for splicing painted a distinct picture of gene expression heterogeneity across subsets of helper T cells, with relatively uniform expression of many genes giving way to highly specific expression patterns based on splicing. Analysis of one such gene, *CRIP1*—encoding an intracellular zinc transport protein typically expressed in effector memory CD4^+^ T cells^13^ is shown on **Fig. 2c,d**. Other relevant genes include *FOS, ANXA1, TCF7, INPP4B*, and *MALAT1* (**Supplementary Figs. 2,3, Supplementary Table 1**).

### SANSARA investigation of T_reg_ subsets

We performed an analysis of the T_reg_ subpopulation of CD4^+^ T cells^23,24^ in order to assess what functionally relevant information could be unearthed with the use of splicing-aware scRNA-Seq analysis. At several resolutions, SANSARA consistently distinguished three major T_reg_ clusters (**Fig. 3a-d**), corresponding to *naïve, activated*, and *effector* T_reg_s, as classified in a recent deep scRNA-Seq investigation^25^. We have retained these cluster designations for consistency. Splicing-unaware T_reg_ clusters (**Fig. 3d**) mapped similarly on the splicing-unaware UMAP (**Fig. 3c**). Corresponding clusters could be also identified in splicing-unaware analysis (**Fig. 3a**), and localized similarly yet not identically within the splicing-aware UMAP (**Fig. 3b**).

**Figure 3.**
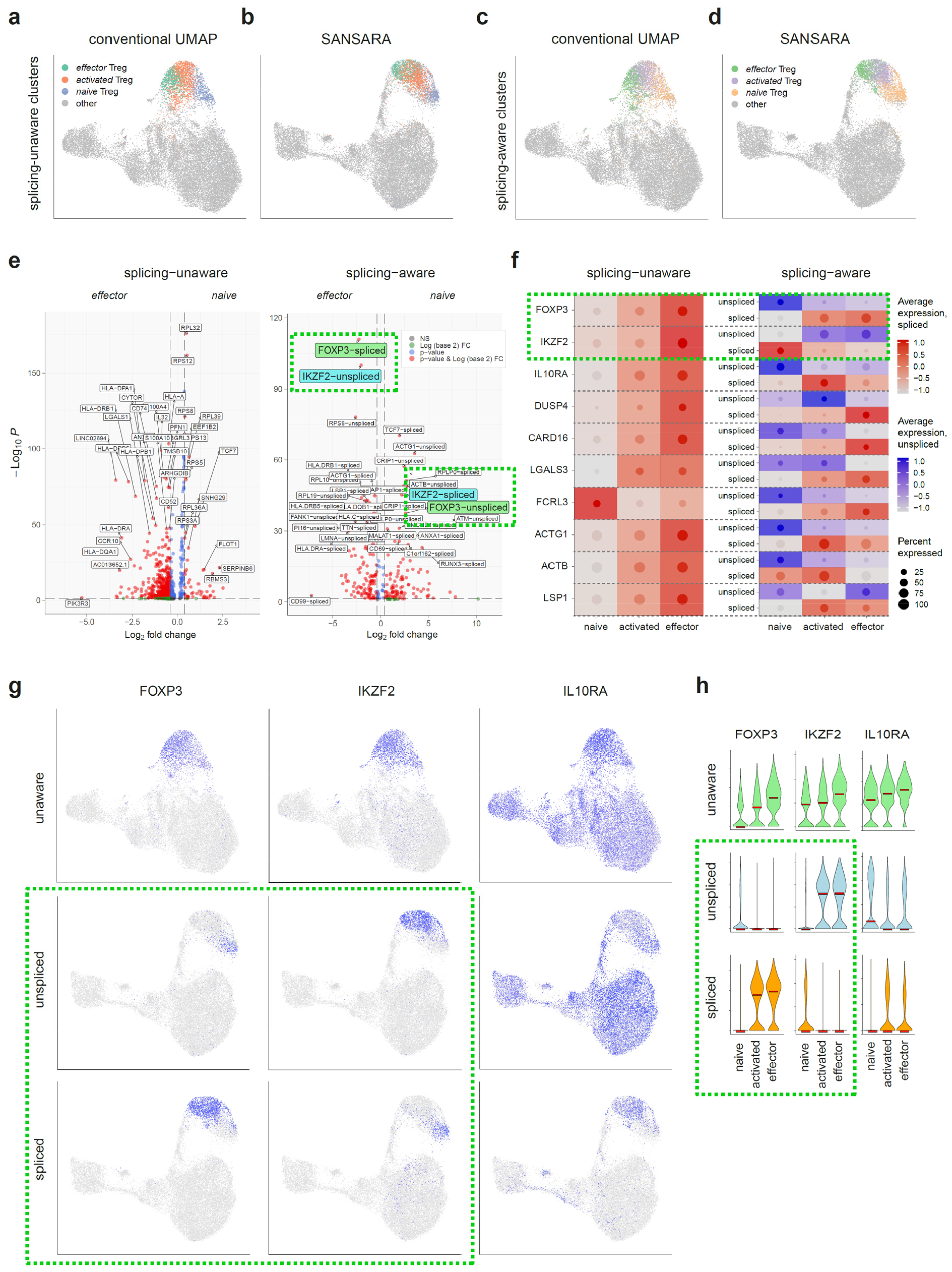
Splicing-aware investigation of T_reg_ heterogeneity. **a-d**. Cross-positioning of splicing-unaware (a, b) and splicing-aware (c, d) naive, activated, and effector T_reg_ clusters in splicing-unaware (a, c) and splicing-aware (b, d) datasets. Splicing-unaware *activated* T_reg_ cluster (a,b) is a product of merging of the two corresponding clusters, see **Supplementary Fig. 1c**, resolution 2.0). **e**. Volcano plots of differentially-expressed genes between *naive* and *effector* T_reg_ clusters from splicing-unaware (left) and -aware (right) datasets. **f**. Dot plot of standardized scaled expression of selected genes in three T_reg_ clusters in splicing-unaware (left) and -aware (right) datasets. Diameter of the dot shows proportion of cells expressing the gene. Background heatmap color corresponds to the color of the dot and reflects average expression of unaware or spliced (red) or unspliced (blue) gene forms. **g**. Splicing-unaware (top) and SANSARA (middle, bottom) UMAP plots showing expression of *FOXP3, IKZF2*, and *IL10RA*. **h**. Violin plots showing conventional splicing-unaware (top) and splicing-adjusted (middle, bottom) *FOXP3, IKZF2*, and *IL10RA* expression across three T_reg_ clusters. Dashed green rectangles highlight expression of spliced and unspliced *FOXP3* and *IKZF2*.

Many of the revealed differences in expression of spliced versus unspliced transcripts were unexpected and informative, and could thus meaningfully shape our understanding of the underlying functional state of different T_reg_ subsets (**Fig. 3e-h, Supplementary Figs. 2-4, Supplementary Table 2**).

In particular, the gene encoding the T_reg_ master transcription factor *FOXP3*^26,27^, was mostly expressed in the unspliced form in *naïve* T_reg_s, presumably reflecting their readiness yet not involvement in active regulatory functions. In *activated* and *effector* T_reg_s, *FOXP3* was mostly expressed in a spliced form. In contrast, another T_reg_-characteristic transcription factor, IKZF2 (Helios) was expressed in the unspliced form in *activated* and *effector* Tregs, while *naïve* Tregs predominantly contained spliced *IKZF2* mRNA (**Fig. 3e-h**). Previous data from mouse models have shown that the Helios transcription factor ensures T_reg_ survival and lineage stability through activation of the IL-2Rα–STAT5 pathway and STAT5-dependent stabilization of *FOXP3* expression^28,29^. Our data indicate that the interplay between these two transcription factors may be more complex at the level of splicing regulation. Furthermore, activated and effector T_reg_s were respectively characterized by expression of unspliced and spliced forms of *DUSP4* (dual-specificity phosphatase-4) (**Fig. 3f**), which encodes a protein that is involved with the regulation of STAT5 protein stability^30^.

All CD4^+^ T cells expressed *IL10RA* according to conventional GEX analysis, but SANSARA revealed that the spliced form of *IL10RA* was almost exclusively observed in *activated* and *effector* T_reg_s. Unspliced *IL10RA* expression was more prominent in *naïve* T_reg_s and non-T_reg_ CD4^+^ T cells (**Fig. 3f-h**). Expression of IL10RA on T_reg_s is important for a feed-forward loop in which IL-10RA signaling reinforces IL-10 secretion by T_reg_s, critical for proper control of Th17 subset activity^31-32^.

We also observed a number of other cluster-specific patterns of splicing behavior. The spliced form of *LGALS3* (encoding galectin-3) was predominantly present in the *effector* T_reg_ cluster, while the unspliced form was present in *naïve* and *activated* T_reg_s (**Fig. 3f, Supplementary Fig. 4**). Galectin-3 has been shown to regulate T_reg_ frequency and function in mouse models of *Leishmania major* infection^33^ and autoimmune encephalomyelitis^34^. Reports have also shown that *LGALS3* expression is increased in human T_reg_s through a transcriptional mechanism involving the ubiquitin D (*UBD*) gene, which is downstream element of FOXP3^35^.

The *activated* T_reg_ cluster was previously shown to express increased levels of *FCRL3* gene encoding Fc receptor-like protein 3^25^. FCRL3 receptor stimulation of T_reg_s has been shown to inhibit their suppressive function and induce IL-17, IL-26, and IFNγ production as well as expression of the Th17-defining transcription factor RORγt^36^. SANSARA revealed that spliced *FCRL3* is mostly expressed in effector T_reg_s, potentially linking FCRL3 to self-restraint of effector T_reg_ function (**Fig. 3f**).

*Naïve* T_reg_s preferentially expressed unspliced transcripts of the cytoskeleton-related protein genes *ACTG1* and *ACTB*^22^, whereas expression of the spliced forms of these transcripts was more characteristic of *activated* T_reg_s (**Fig. 3f, Supplementary Figs. 2,4**). The spliced form of the *LSP1*, which encodes leukocyte-specific protein 1, potentially associated with negative regulation of T cell migration^37^, was mostly detected in the *activated* T_reg_ cluster (**Fig. 3f, Supplementary Fig. 4**).

The *naïve* T_reg_ cluster was also characterized by expression of spliced *TCF7* (a marker of T cells with high capacity for self-renewal^17^), *SKAP1* (an immune cell adaptor that regulates T-cell adhesion and optimal cell growth^38^), *RBMS1* (encodes RNA-binding motif 1, a single-stranded-interacting protein involved in helper T cell and T_reg_ post-transcriptional gene regulation^39^), *PTGER2* (encodes PGE2 receptor EP2, involved in differentiation and expansion of helper T cell subsets^40^), and *MALAT1* (a long noncoding RNA linked to regulation of helper T cell differentiation^21^) (**Fig. 3f, Supplementary Figs. 2,4, Supplementary Table 2**).

### SANSARA investigation of Th1/cytotoxic CD4+ subsets

Next, we focused on analyzing the heterogeneity of gene splicing states in Th1 and cytotoxic CD4^+^ subsets. In SANSARA analysis, a number of genes characteristic for cytotoxic lymphocytes showed heterogeneous splicing behavior across the clusters, including *NKG7, PRF1, GNLY, GZMA*^41,42^, *CCL5, FGFBP2, CST7*^43^, *ADGRG1* (*GPR56*)^44^, *PLEK*^45^, transcription factors *HOPX*^46^ and *ETS1*^47^ (**Supplementary Figs. 5-7**).

For example, although we detected *PRF1* expression in T_reg_, *Eff-Mem Th22, Eff-Mem Th1*, and *Temra cytotoxic Th1* clusters with conventional splicing-unaware analysis, SANSARA revealed that the spliced form of this gene is almost exclusively expressed in the *Temra cytotoxic Th1* cluster (**Supplementary Fig. 6**).

Further partitioning of the *Temra cytotoxic Th1* cluster based on splicing of *GNLY* may be indicative of heterogeneous cytotoxic functions performed by distinct subpopulations of helper T cells (**Supplementary Fig. 7**).

*CCL5*, which encodes a characteristic cytokine of cytotoxic lymphocytes, was mostly present in the unspliced form, except within a compact subpopulation within *Eff-Mem Th1* cluster, suggesting that circulating Th1 and cytotoxic CD4^+^ T cells exist in a relatively dormant state, but remain ready to rapidly produce CCL5 in case of activation^48^ (**Supplementary Fig. 5**).

*HOPX*—encoding the transcription factor which is thought to be involved in imprinting for terminal effector differentiation^46,47^, was uniformly expressed in *Eff-Mem Th1* and *Temra cytotoxic Th1* clusters in splicing-unaware analysis, but SANSARA revealed distinctive expression patterns for its spliced versus unspliced forms (**Supplementary Fig. 6**).

Another transcription factor, ETS1 (which is involved in Th1 differentiation and IFNγ production^47^, was uniformly expressed across CD4^+^ T cells in splicing-unaware analysis. SANSARA showed that spliced *ETS1* is mostly expressed in a compact subpopulation within the *Temra cytotoxic Th1* cluster (**Supplementary Fig. 7**).

SANSARA also revealed that a compact subset within the Temra *cytotoxic Th1* cluster is characterized by spliced *ADGRG1*, which encodes a GPR56 protein linked to extracellular signaling and was established as a marker of IFNγ- and TNF-producing Th1 cells^44^ (**Supplementary Fig. 6**).

## Discussion

The ability to profile single-cell transcriptomes has fundamentally changed our approach to studying the diversity, combinations, and functional impact of genetic programs in living cells^49,50^. However, the functional implementation of genetic programs occurs at multiple levels, not just at the level of the quantity of produced and stored RNA. Ideally, analyzing the transcriptomes of single cells could also reveal the proportion of spliced RNA molecules, which directly affect the functional activity of both mRNAs and non-coding RNAs, as well as offer the insights into alternative splicing^5,51^ and trans-splicing^52^.

However, this has proven methodologically challenging, as the use of either 5’ or 3’ end-labeling of RNA molecules with molecular barcodes—alongside inherent limitations of high-throughput sequencing methods—have restricted our ability to comprehensively derive such information for a given RNA molecule^3,51^.

Algorithms developed by the Kharchenko and Yosef teams^6,10^ have enabled estimation the RNA processing velocity, making it possible to study transitions between cell types as they differentiate and change gene expression programs at the post-analysis level of scRNA-Seq data. In SANSARA, we have exploited these same algorithms to transform splicing-unaware gene expression data into a splicing-aware format referred to as the saGEX matrix.

SANSARA operates on information about genes predominantly represented in spliced or unspliced form in a given cell, and can be used to build an alternative UMAP data representation that reveals splicing-aware cell clusters. Obtained saGEX matrices are directly usable for Seurat dimensionality reduction and clustering analysis, allowing for seamless transition from conventional scRNA-Seq data analysis. Based on the obtained results, we believe that we managed to find a non-disruptive way to exploit splicing information in scRNA-Seq clustering and dimensionality reduction. Resulting UMAP topology and cluster annotations closely resemble the results of the conventional analysis, and offer intuitively understandable, and easy-to-implement analytical approach.

The differentiation between spliced and unspliced mRNA enabled by SANSARA facilitates discovery of distinct features that are informative about cell subset heterogeneity. As a demonstration, we have applied SANSARA to peripheral CD4^+^ T cell scRNA-Seq data, revealing several unexpected features in different helper T cell subsets. In T_reg_s, we uncovered reciprocal splicing interplay between the master transcription factors FoxP3 and Helios, alongside exclusive expression of the spliced form of *IL10RA* in activated and effector T_reg_s. These findings have significant implications for our understanding of T_reg_ biology^53^ and T_reg_-based therapy developments^54^. Investigation of Th1 and cytotoxic CD4^+^ T cells also revealed numerous unexpected splicing-related heterogeneities, indicating a diverse composition of heterogeneous helper T cell functions associated with type 1 immune response.

Based on these demonstrations, we believe that SANSARA could change the way we analyze single cell transcriptomic data, opening up a new—and currently unexploited— dimension for investigating the critically important role of splicing regulation in cellular gene expression programs.

## Methods

### saGEX matrix calculation

Raw scRNA-seq data were mapped to the genome using cellranger (v7.1) *count*, taking into account intronic sequences^55^. Subsequently, the velocyto utility^6^ was used to count UMIs belonging to unspliced and spliced forms of RNA; cDNAs containing at least some intronic sequences were classified as unspliced, while remaining cDNA reads were identified as spliced. Using the veloVI (v.0.3.0) package^10^, we selected highly variable genes and genes with a sufficient number of unspliced and spliced forms for further analysis. For the selected genes, phase portraits reflecting the balance of spliced and unspliced forms were constructed. The splicing score was calculated for each gene in each cell based on the gene-specific phase portrait. The normalized expression of variable genes was then multiplied by the splicing score value of each gene in each cell. We divided the resulting metric into spliced and unspliced—negative values were defined as the “expression” of the spliced form of the gene, while positive values described the unspliced form of the gene—and took the modulo values. The resulting splicing-aware gene expression (saGEX) matrix of spliced/unspliced counts was used for downstream Seurat (v.5.0.1) normalization, dimensional reduction and clustering^56^.

### Integration and clustering

The Harmony pipeline was used for the integration separately for the GEX and saGEX datasets^12^. These datasets were independently normalized using the `LogNormalise` function and integration features were selected with the `SelectIntegrationFeatures` function in Seurat. After merging the datasets, variable features of the merged object were set to selected integrated features. Principal Components (PCs) were calculated from scaled integration features. Harmony was run with the default options, and the top 25 corrected Harmony PCs were used to generate UMAP plots. Clustering analysis was performed on Harmony PCs via the `FindNeighbours` and `FindClusters` Seurat functions. Clustering trees were built using the ‘clustree’ R package for the set of resolutions from 0 to 2.5.

### Differential expression and annotation

For differential expression analysis, the `FindMarkers` and `FindAllMarkers` Seurat functions were used. As these datasets were previously characterized, annotation was performed on the basis of the extensive reference^11^, the composition of clusters at resolution level 2.0, and the differential expression results. If cells from the same proposed annotation belonged to several clusters, these clusters were merged. Dotplots and volcano plots were generated via the `DotPlot` Seurat function and EnhancedVolcano R package.

## Data availability

Raw scRNA-seq data can be accessed on NCBI Sequence Read Archive under the BioProject PRJNA995237 accession. Tables with saGEX and GEX values as well as our custom code pipeline are available at Github (https://github.com/EvgenEgorov/SANSARA).

---

**Supplementary Fig. 1.**
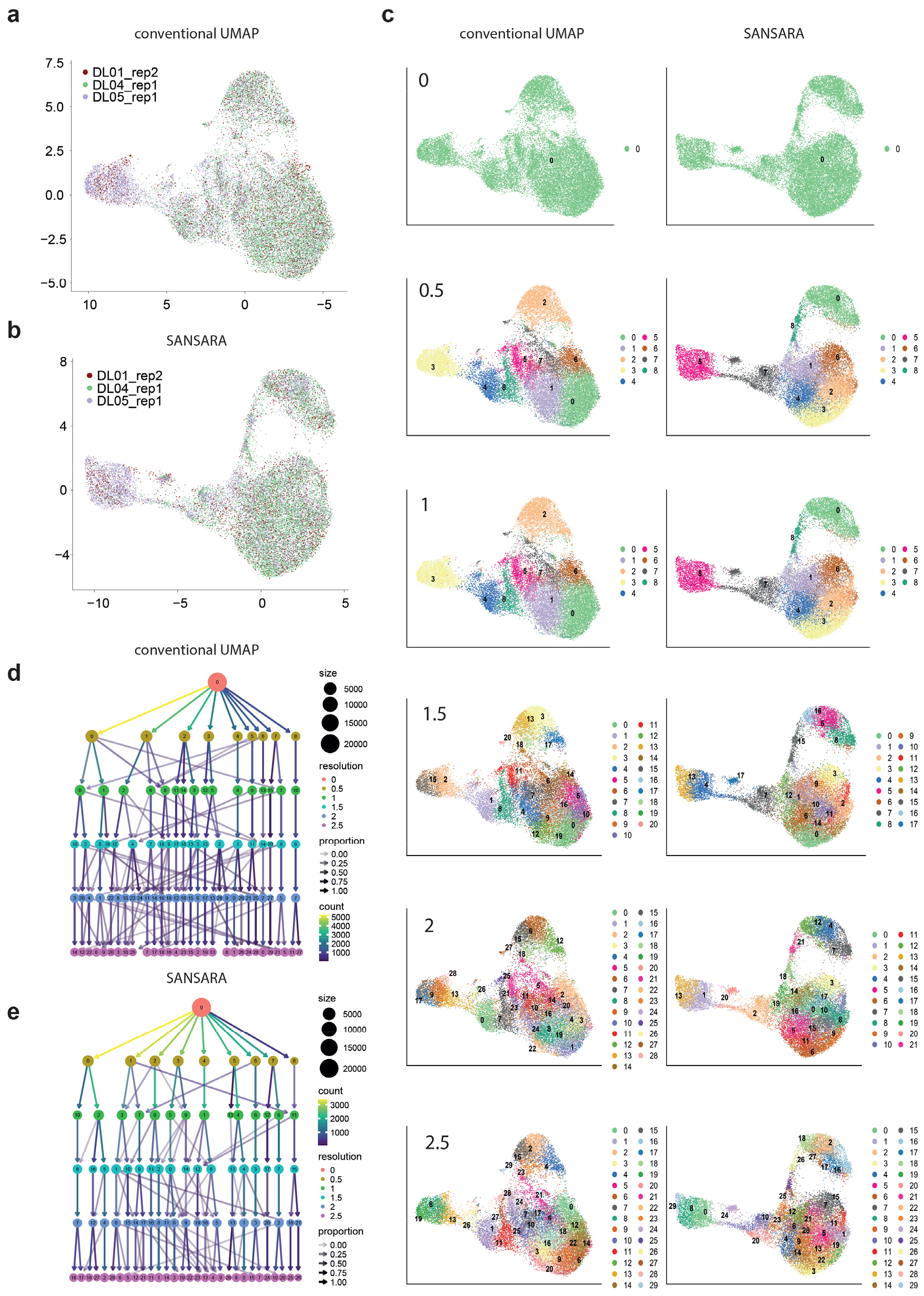
Integration, cluster stability. **a,b**. Harmony integration of scRNA-Seq data for the three donors performed with conventional (a) and splicing-aware (b) datasets. **c**. Clustering at different UMAP resolutions. **d,e**. Clustering trees for splicing-unaware (d) and splicing-aware (e) datasets.

**Supplementary Figure 2.**
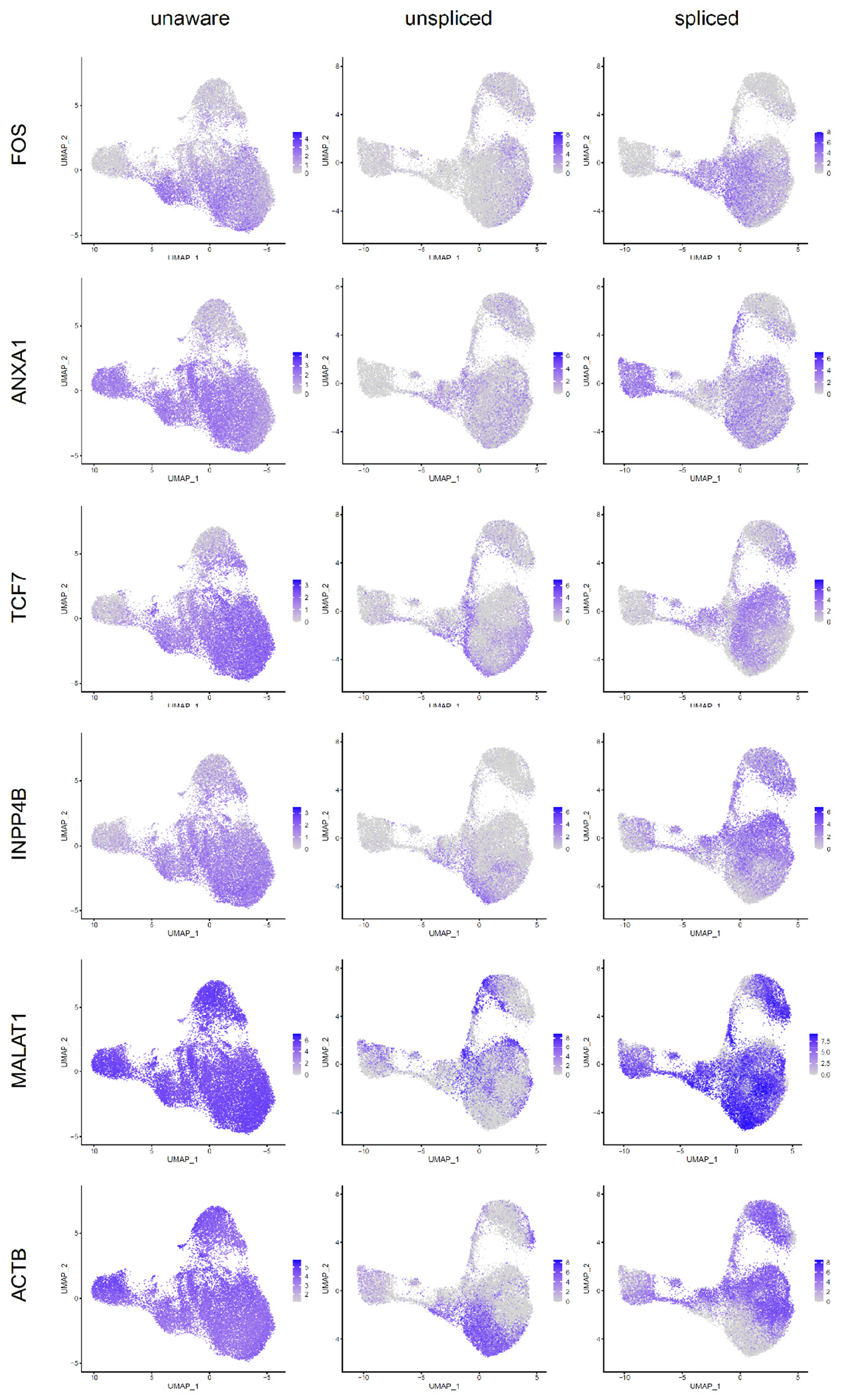
Selected genes characterized by heterogeneous expression of spliced and unspliced forms. Splicing-unaware UMAP plots are shown at left, center and right panels show splicing-aware UMAP plots. *FOS*—encoding a c-Fos protein which interacts with c-Jun, forming heterodimeric AP-1 transcription factor that prominently affects CD4^+^ T cell differentiation^14^. *ANXA1*—encoding Annexin A1, the key driver of glucocorticoid anti-inflammatory effects, involved in T-cell differentiation, altering the strength of TCR signaling^15^ and Th1-Th2 counterbalance driven by GATA3 and TBX21 transcription factors^16^. *TCF7*—encoding transcription factor T cell factor 1 which marks CD4+ T cells ability to self-renew^17^ and which expression goes down along with effector T cell differentiation^18^, especially towards CD4+ cytotoxic T cells^19^. *INPP4B*—encoding inositol poly-phosphate 4-phosphatase that was suggested to play role in T cell proliferation, survival and differentiation^20^. *MALAT1*—long noncoding RNA, reported as regulator of helper T cell differentiation from naïve CD4+ T cells^21^. *ACTG1* and *ACTB*—cytoskeleton-related protein genes^22^.

**Supplementary Figure 3.**
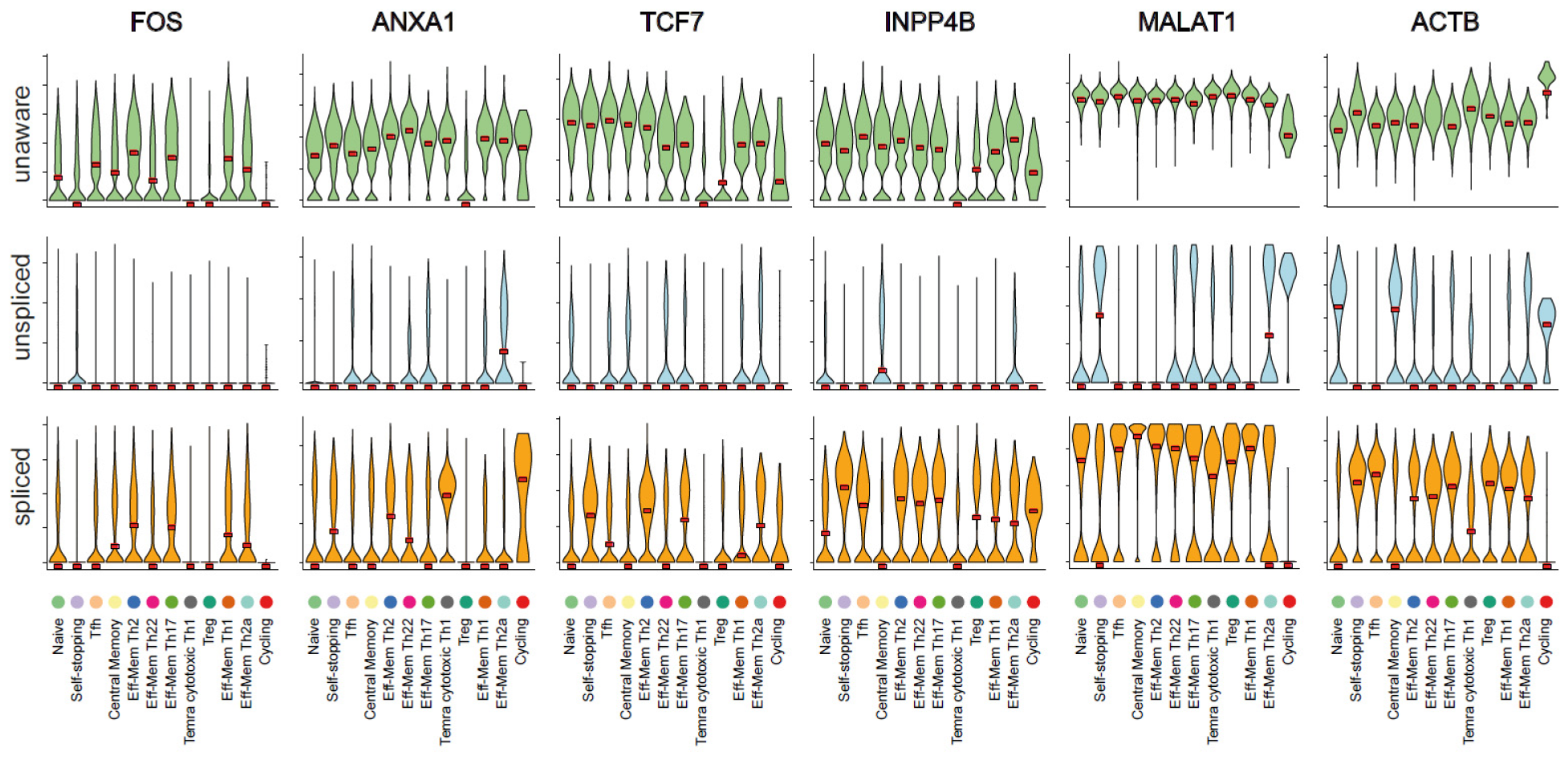
Selected genes characterized by heterogeneous expression of spliced and unspliced forms. Violin plots of splicing-unaware (top) and -aware (middle, bottom) gene expression across clusters are shown. *FOS*—encoding a c-Fos protein which interacts with c-Jun, forming heterodimeric AP-1 transcription factor that prominently affects CD4^+^ T cell differentiation^14^. *ANXA1*—encoding Annexin A1, the key driver of glucocorticoid anti-inflammatory effects, involved in T-cell differentiation, altering the strength of TCR signaling^15^ and Th1-Th2 counterbalance driven by GATA3 and TBX21 transcription factors^16^. *TCF7*—encoding transcription factor T cell factor 1 which marks CD4+ T cells ability to self-renew^17^ and which expression goes down along with effector T cell differentiation^18^, especially towards CD4+ cytotoxic T cells^19^. *INPP4B*—encoding inositol poly-phosphate 4-phosphatase that was suggested to play role in T cell proliferation, survival and differentiation^20^. *MALAT1*—long noncoding RNA, reported as regulator of helper T cell differentiation from naïve CD4+ T cells^21^. *ACTG1* and *ACTB*—cytoskeleton-related protein genes^22^.

**Supplementary Figure 4.**
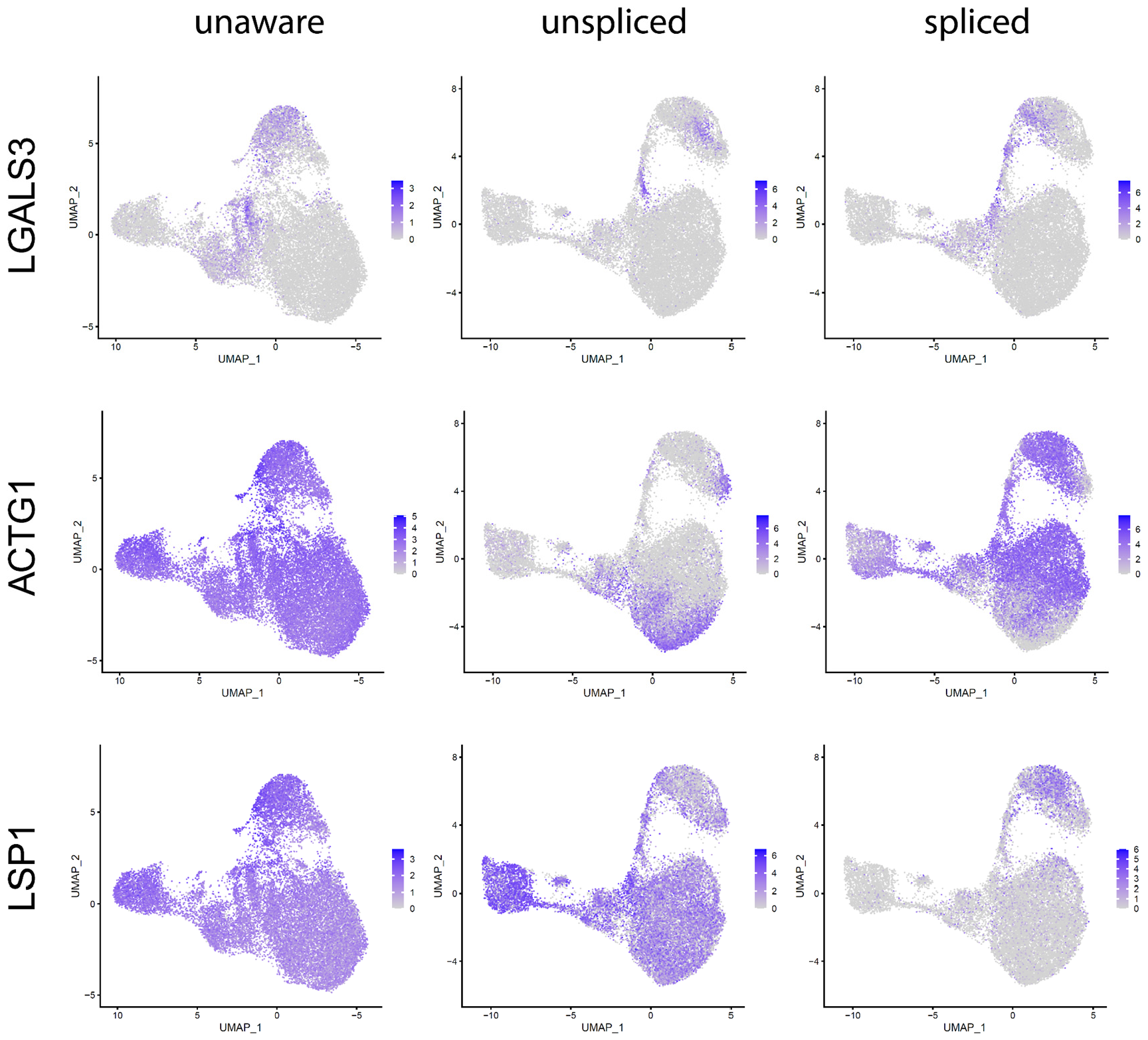
Selected genes characterized by heterogeneous expression of spliced and unspliced forms in T_reg_ clusters. The lefthand column shows splicing-unaware UMAP plots, center and righthand columns show splicing-aware UMAP plots.

**Supplementary Figure 5.**
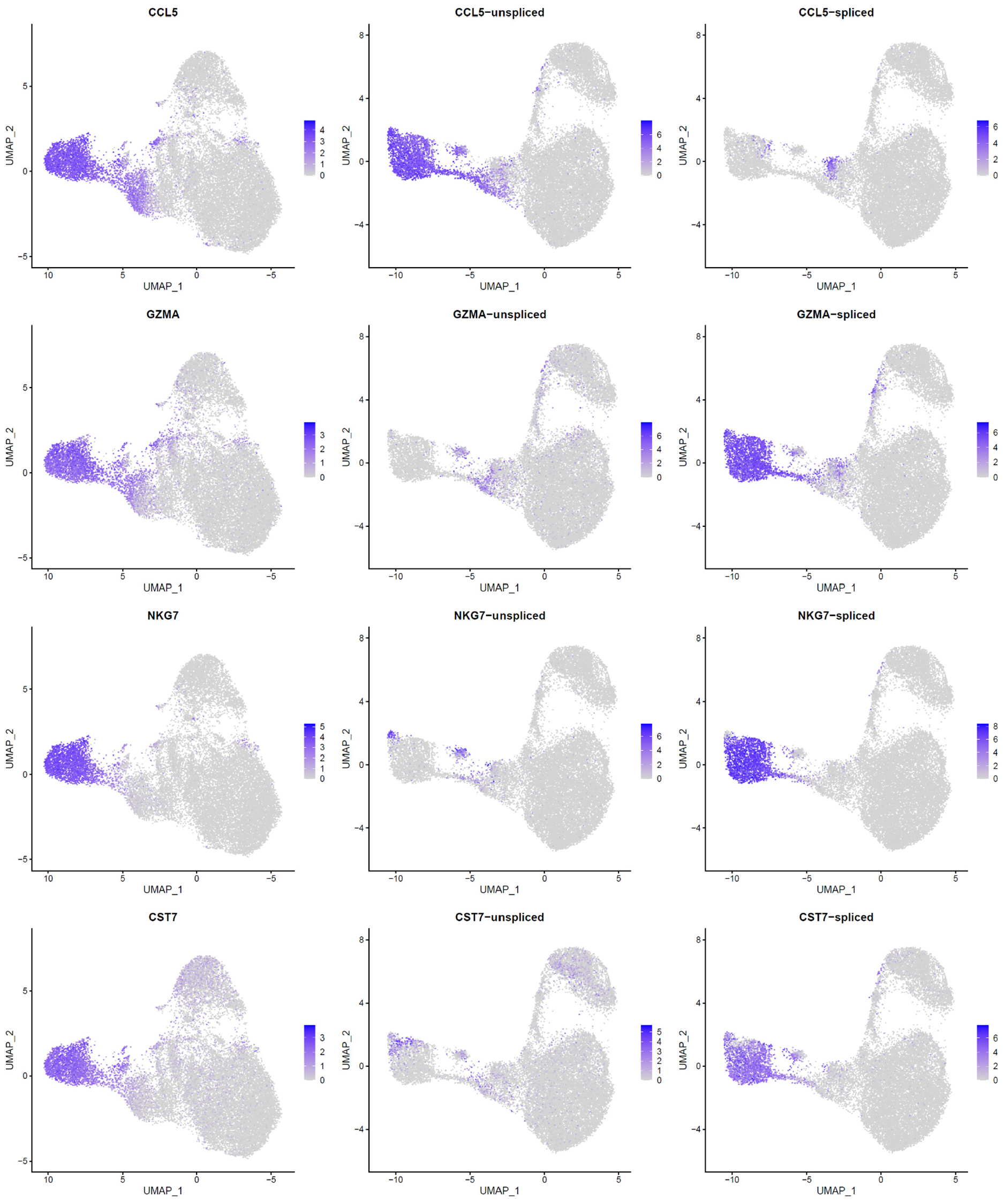
Heterogeneous expression of spliced and unspliced forms of *CCL5, GZMA, NKG7*, and *CST7*. Lefthand column shows splicing-unaware UMAP plots for comparison purposes.

**Supplementary Figure 6.**
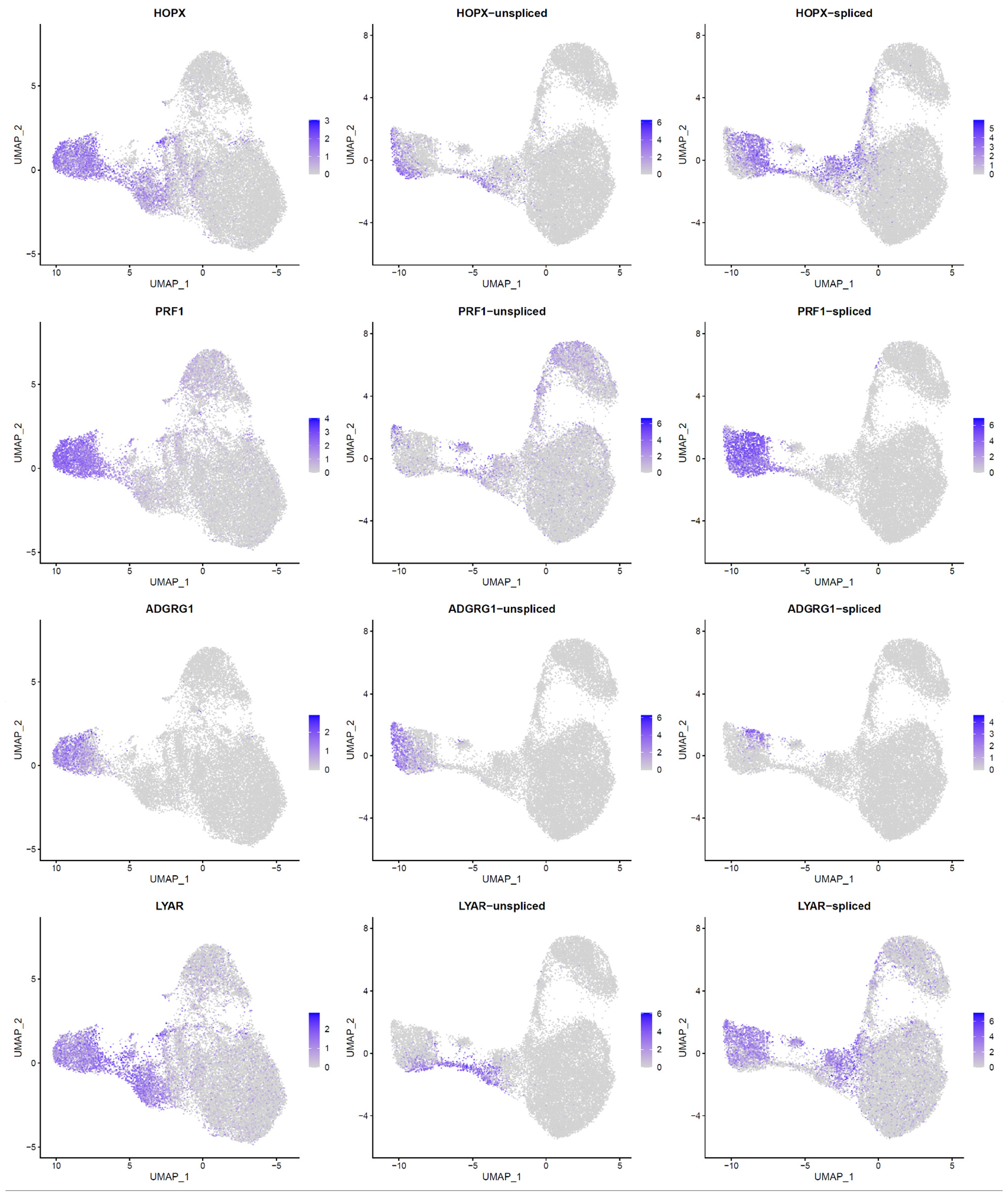
Heterogeneous expression of spliced and unspliced forms of *HOPX, PRF1, ADGRG1*, and *LYAR*. Lefthand column shows splicing-unaware UMAP plots for comparison purposes.

**Supplementary Figure 7.**
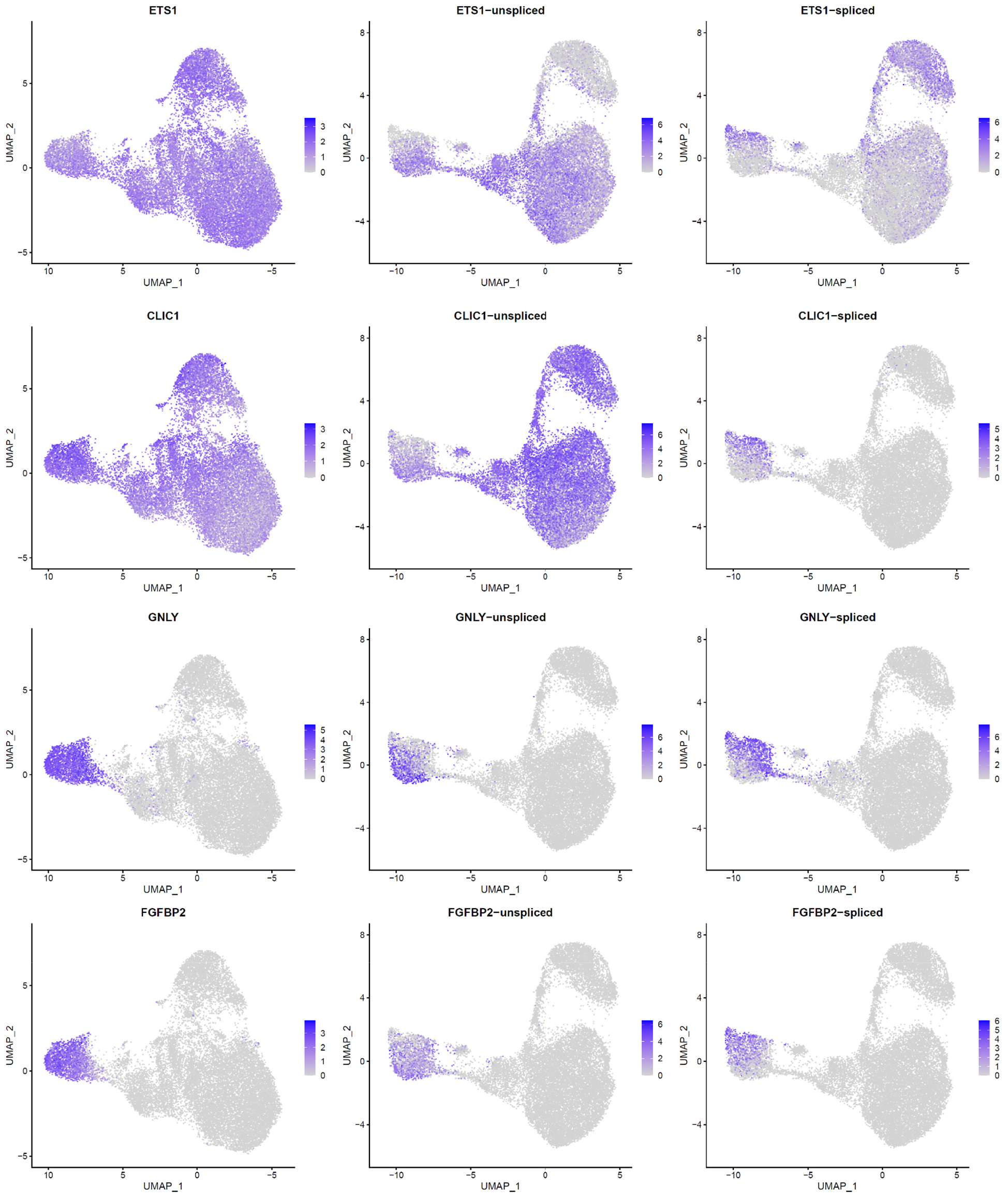
Heterogeneous expression of spliced and unspliced forms of *ETS1, CLIC1, GNLY*, and *FGFBP2*. Lefthand column shows splicing-unaware UMAP plots for comparison purposes.

